# A new fungal taurine biosynthetic pathway promotes metabolic fitness and virulence in *Candida albicans*

**DOI:** 10.64898/2026.08.27.747329

**Authors:** Anagha C.T. Menon, Faiza Tebbji, Nathan Ghafari, Lekha Sleno, Adnane Sellam

**Affiliations:** Montreal Heart Institute, Université de Montréal, Montréal, QC, Canada; Department of Microbiology, Infectious Diseases and Immunology, Faculty of Medicine, Université de Montréal, Montréal, QC, Canada; Chemistry Department, University of Quebec in Montreal (UQAM), Montreal, QC, Canada

## Abstract

Taurine is an abundant sulfur-containing metabolite with diverse roles in cellular physiology across many organisms, yet its biosynthesis and biological functions remain largely unexplored in fungi. Here, we provide evidence for endogenous taurine production in the major human fungal pathogen *Candida albicans* and identify Csd1, a cysteine sulfinic acid decarboxylase (CSAD)-related protein, as a major determinant of this process. Loss of *CSD1* nearly abolished intracellular taurine and caused extensive remodeling of sulfur metabolism, including cysteine accumulation and altered abundance of methionine-cycle metabolites. Consistent with these metabolic defects, *csd1* cells exhibited impaired growth and increased sensitivity to cysteine, oxidative and osmotic stresses, elevated temperature, reactive sulfur species, and the antifungal drugs amphotericin B and caspofungin. Exogenous taurine selectively rescued a subset of these phenotypes, indicating that *CSD1* loss causes both taurine-dependent and broader metabolic defects. Csd1 was also required for normal hyphal morphogenesis, and *csd1* cells displayed markedly attenuated virulence in a *Galleria mellonella* systemic infection model. Comparative sequence analysis revealed conservation of key features of the pyridoxal 5′-phosphate-dependent catalytic machinery shared with mammalian and bacterial CSADs, together with divergence within the predicted substrate-recognition pocket. Our genetic data further suggest that taurine production in *C. albicans* differs from the canonical metazoan cysteine sulfinic acid pathway and may involve branched or redundant routes. Together, these findings establish endogenous taurine production as a new facet of fungal sulfur metabolism and identify Csd1-dependent metabolism as an important contributor to sulfur homeostasis, stress adaptation, morphogenesis, and pathogenic fitness in *C. albicans*.

**Author Summary:** Taurine is a sulfur-containing molecule abundant in animals, where it contributes to numerous physiological processes. Although taurine has also been detected in fungi, whether fungi can synthesize it and what biological functions it might serve have remained poorly understood. Here, we show that the highly prevalent human fungal pathogen *Candida albicans* produces taurine and identify Csd1 as a protein required for efficient taurine production. Cells lacking *CSD1* contained almost no taurine and showed broad alterations in sulfur metabolism, including cysteine accumulation. These cells grew poorly and were more vulnerable to several environmental stresses and antifungal drugs. They were also defective in forming hyphae, a growth form associated with *C. albicans* pathogenicity, and were markedly less virulent in an infection model. Interestingly, supplying taurine restored some, but not all, of these defects, indicating that loss of *CSD1* affects both taurine production and broader sulfur metabolic homeostasis. Our findings establish taurine biosynthesis as a new facet of fungal sulfur metabolism and reveal its importance for the ability of *C. albicans* to adapt to stressful conditions and maintain pathogenic fitness.

## Introduction

Fungal infections represent an escalating global health threat, causing an estimated 3.8 million deaths annually [1, 2]. Recognizing this burden, the World Health Organization (WHO) established the Fungal Priority Pathogens List in 2022, placing the opportunistic yeast *Candida albicans* among the highest-priority fungal pathogens because of rising antifungal resistance, limited therapeutics, and persistently high mortality [2–6]. More recently, the WHO launched the 2026 “Blueprint for Strengthening Responses to Fungal Disease and Antifungal Resistance”, which identifies research and innovation as a central priority to accelerate the discovery of new antifungal targets and therapeutic strategies [7]. Despite currently available antifungal therapies, systemic candidiasis remains associated with mortality rates exceeding 50% [2, 8], while treatment continues to rely on a limited repertoire of antifungal classes and molecular targets. The eukaryotic nature of fungi further constrains antifungal drug development, as many cellular processes are shared with their human hosts, limiting opportunities for selective therapeutic intervention. Together, these limitations underscore the urgent need to uncover fungal-specific vulnerabilities that can be exploited for antifungal drug development.

Sulfur is available within the host in a chemically diverse array of inorganic and organic compounds, including sulfate, sulfite, cysteine, methionine, glutathione, and sulfonates [9]. We recently showed that *C. albicans* possesses an extensive sulfur-utilization network that enables the fungus to exploit alternative sulfur sources, including glutathione and sulfonates such as taurine [10]. Taurine is a highly abundant sulfur-containing metabolite in mammalian tissues and biological fluids, where it participates in osmoregulation, membrane stabilization, redox homeostasis, bile acid conjugation, and cellular stress responses [11–13]. Beyond free taurine, more than 100 taurine-conjugated metabolites and taurine derivatives have been reported in the gut and circulation, representing a remarkably diverse pool of sulfur-containing compounds that *C. albicans* could potentially exploit [14]. Although microorganisms can utilize taurine as a sulfur source through desulfonation pathways [10, 15], the biosynthesis and physiological functions of endogenous taurine remain poorly understood across the microbial world, including fungi.

In metazoans and algae, taurine is synthesized primarily from cysteine through the cysteine sulfinic acid (CSA) pathway [16, 17]. Cysteine is first oxidized to CSA by cysteine dioxygenase [18] and subsequently decarboxylated to hypotaurine predominantly by cysteine sulfinic acid decarboxylase (CSAD) [16], with glutamate decarboxylase (GAD) contributing to a lesser extent [19, 20]. Hypotaurine is then converted to taurine either enzymatically by hypotaurine dehydrogenase or through spontaneous oxidation [11]. An alternative route can proceed through the oxidation of CSA to cysteic acid, followed by its direct decarboxylation to taurine [21]. Taurine biosynthesis has also been demonstrated in bacteria, including cyanobacteria [22, 23]. Evidence for endogenous taurine biosynthesis has similarly emerged in fungi, with taurine and hypotaurine detected in *Yarrowia lipolytica* and *Geotrichum candidum* and components of the CSA pathway conserved across several fungal species [24, 25]. However, the genetic architecture of taurine biosynthesis and the physiological functions of endogenously produced taurine remain largely unresolved in fungi. This knowledge gap is particularly relevant to *C. albicans*, which encounters taurine in host environments and possesses an expanded repertoire of desulfonating enzymes capable of mobilizing taurine-derived sulfur.

Here, we provide functional evidence for endogenous taurine biosynthesis in *C. albicans* and identify Csd1 as a major determinant of taurine production. Loss of *CSD1* markedly reduced intracellular taurine, disrupted sulfur amino acid homeostasis, and compromised growth, stress adaptation, hyphal morphogenesis, and virulence. Together, our findings establish taurine biosynthesis as a new facet of fungal sulfur metabolism and link this pathway to sulfur homeostasis, stress adaptation, and pathogenic fitness in *C. albicans*.

## Results

### Evidence of taurine biosynthesis in *C. albicans*

Our previous targeted metabolomic profiling revealed increased taurine and hypotaurine abundance in *C. albicans* cells exposed to hypoxia compared with normoxic conditions (**Figure S1**) [26]. Because these experiments were performed in YPD, a complex medium containing animal-derived peptone, we considered the possibility that the detected taurine could have originated from the medium rather than from fungal metabolism. To exclude this potential source, *C. albicans* clinical isolate SC5314 and two laboratory-derived strains (CAI4 and SN250) were cultivated in a chemically defined minimal dextrose medium (MDM), and taurine levels were quantified by liquid chromatography-tandem mass spectrometry (LC-MS/MS). Taurine was readily detected in extracts from MDM-grown cells, supporting its endogenous production by *C. albicans* (**Figure 1A**). At a cell density of 10^6^ cells, taurine concentrations in fungal cells were in the micromolar range (1.5-2.2 µM).

**Figure 1.**
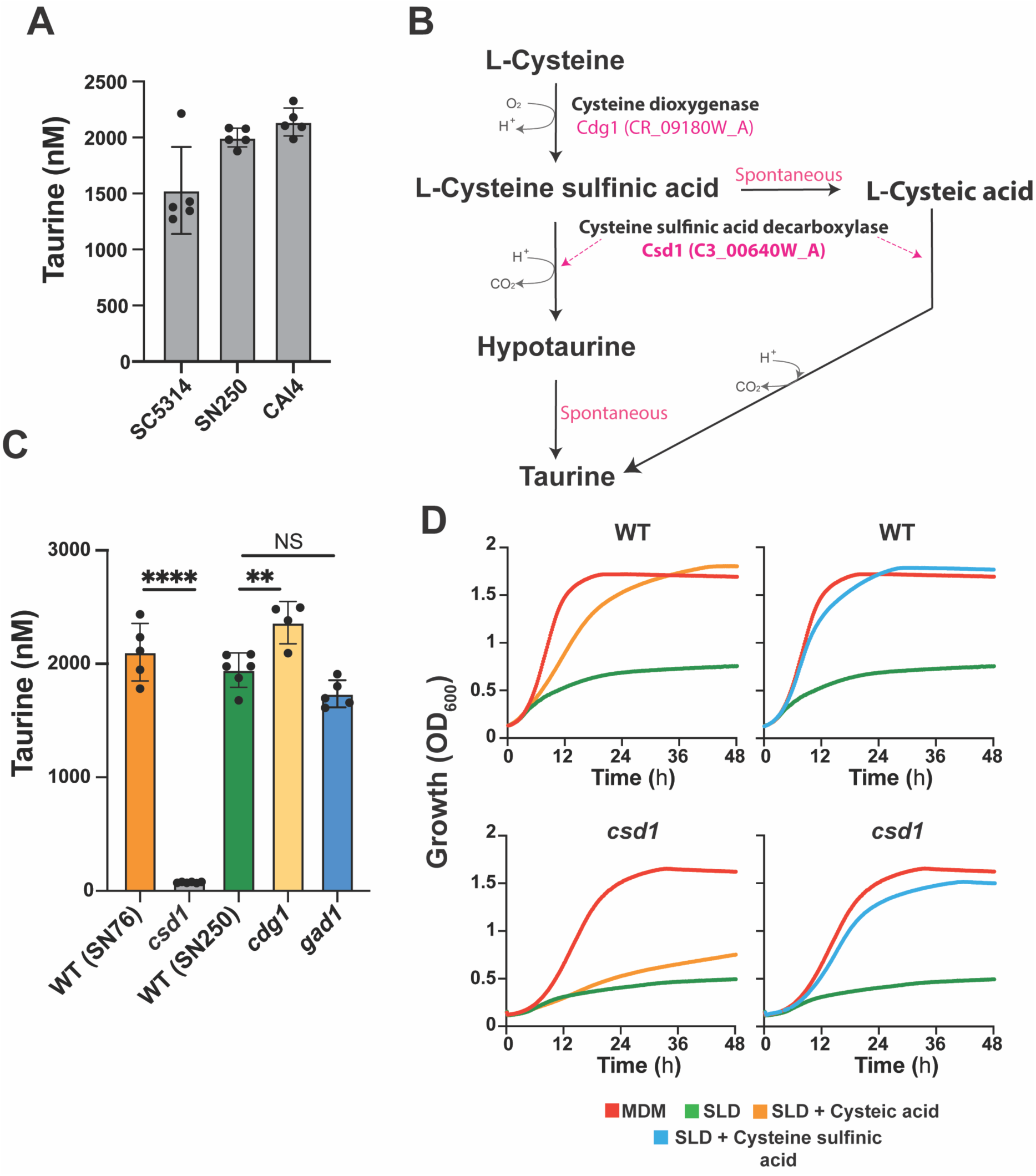
Taurine production in *C. albicans* requires Csd1. (**A**) Taurine levels in the *C. albicans* clinical isolate SC5314 and laboratory strains SN250 and CAI4 grown in chemically defined minimal dextrose medium (MDM). Taurine was quantified by LC-MS/MS in extracts prepared from 10^6^ cells. (**B**) Proposed routes potentially contributing to taurine production in *C. albicans*, based on pathways characterized in metazoans and algae. Cysteine is converted to cysteine sulfinic acid, which can be decarboxylated to hypotaurine and subsequently oxidized to taurine, or oxidized to cysteic acid and subsequently decarboxylated to taurine. Candidate *C. albicans* enzymes potentially involved in each step are indicated. (**C**) Intracellular taurine levels in *csd1*, *cdg1*, and *gad1* mutants and their corresponding parental strains. (**D**) Growth of WT and *csd1* cells in SLD supplemented with cysteic acid or cysteine sulfinic acid as the sole sulfur source. Loss of *CSD1* impaired growth on cysteic acid but not on cysteine sulfinic acid. Data in **A** and **C** are presented as means from five independent biological replicates. NS, not significant; **P < 0.01; ****P < 0.0001.

### *CSD1* (C3_00640W) encodes a CSAD-related enzyme required for taurine production

To identify genes involved in taurine biosynthesis, we searched the *C. albicans* genome for homologs of enzymes associated with the canonical CSA pathway. The *C. albicans* genome encodes orthologs of cysteine dioxygenase (Cdg1), glutamate decarboxylase (Gad1), and a putative cysteine sulfinic acid decarboxylase (C3_00640W), suggesting that this organism possesses the enzymatic machinery potentially required for taurine biosynthesis (**Figure 1B**). To determine the contribution of these genes to taurine production, we quantified intracellular taurine levels in the corresponding deletion mutants. Deletion of *GAD1* had no significant effect on taurine levels, while deletion of *CDG1* resulted in a modest increase relative to the congenic parental strains (**Figure 1C**). In contrast, deletion of C3_00640W reduced taurine to levels near the detection limit (**Figure 1C**). Together, these findings identify C3_00640W as a major determinant of taurine production and are consistent with a role for this protein in the taurine biosynthetic pathway of *C. albicans*.

C3_00640W was required for growth on cysteic acid as the sole sulfur source but was dispensable for growth on cysteine sulfinic acid (**Figure 1D**). This differential requirement suggests that cysteic acid utilization involves a C3_00640W-dependent step, potentially its conversion to taurine, consistent with pathways described in bacteria and fungi [27, 28], whereas cysteine sulfinic acid can enter sulfur metabolism through an alternative C3_00640W-independent route. Together, these findings identify C3_00640W as a major contributor of taurine biosynthesis and are consistent with a cysteine sulfinic acid decarboxylase function in *C. albicans*. Based on its homology to mammalian cysteine sulfinic acid decarboxylase (CSAD), we designated C3_00640W as *CSD1* (<u>C</u>ysteine <u>S</u>ulfinic acid <u>D</u>ecarboxylase 1).

Consistent with its requirement for taurine prodction, Csd1 exhibits substantial conservation with characterized cysteine sulfinic acid decarboxylases (CSADs) [22, 29, 30]. Comparative sequence analysis of *C. albicans* Csd1 with human and mouse CSADs and the experimentally characterized CSAD from *Synechococcus sp.* revealed conservation of key features of the pyridoxal 5’-phosphate (PLP)-dependent catalytic machinery (**Figure S2**) [22]. Most notably, the PLP-binding lysine of human CSAD (K305) is conserved in Csd1 (K309), as is the C190-H191-Y192 active-site loop (C194-H195-Y196 in Csd1). In contrast, residues implicated in cysteine sulfinic acid recognition are more divergent. Whereas mammalian CSADs contain the F-X_19_-S-X-Y substrate-recognition signature and the bacterial *Synechococcus* CSAD retains the corresponding W-X_19_-S-X-Y motif, Csd1 contains Y-X_19_-N-X-H at the equivalent positions [22]. Together, these sequence similarities and conservation of key catalytic residues support Csd1 as a divergent fungal member of the CSAD family, while divergence within the putative substrate-recognition pocket suggests evolutionary specialization of the fungal enzyme.

### Genetic inactivation of *CSD1* disrupts sulfur amino acid homeostasis and impairs growth

Cells of the *csd1* mutant displayed a pronounced growth defect, marked by an extended lag phase and increased doubling-time relative to the WT (**Figure 2A-B**). Supplementation of the growth medium with increasing concentrations of taurine failed to rescue this phenotype (**Figure 2B and S3**), indicating that the growth impairment is not simply attributable to taurine deficiency. We therefore hypothesized that the growth defect might instead result from broader metabolic perturbations caused by disruption of the taurine biosynthetic pathway. To investigate this possibility, we performed untargeted metabolomic profiling of *csd1* mutant and WT cells grown under identical conditions. Principal component analysis (PCA) revealed a clear separation between the metabolic profiles of *csd1* and WT cells in both positive and negative ionization modes, indicating substantial metabolic reprogramming in the mutant (**Figure 2C**). Among the 95 unique metabolites that were differentially abundant, 34 were significantly enriched and 61 were depleted in the *csd1* mutant relative to the WT (**Figure 2C-E and Table S1**). The most enriched metabolites included amino acid and amine derivatives, particularly intermediates of cysteine and methionine metabolism (L-cystathionine, S-adenosyl-L-methionine, and 5′-S-methyl-5′-thioadenosine) (**Figure 2D and Table S1**). This pattern is consistent with disruption of the taurine pathway, as several metabolites linked to upstream cysteine biosynthesis and sulfur amino acid metabolism were significantly increased in the *csd1* mutant. As cysteine was not detected in the untargeted metabolomic dataset, we independently quantified its abundance and found that cysteine levels were significantly increased by approximately 2.5-fold in *csd1* cells compared with the WT (**Figure 2F**). Given that elevated cysteine can inhibit fungal growth [31, 32], its accumulation could contribute to the growth defect of the *csd1* mutant. Consistent with this possibility, *csd1* cells exhibited increased sensitivity to exogenous cysteine compared to the WT (**Figure 2G**). Together, these findings indicate that loss of *CSD1* disrupts sulfur amino acid homeostasis and leads to cysteine accumulation, which may contribute to the impaired growth of the *csd1* mutant.

**Figure 2.**
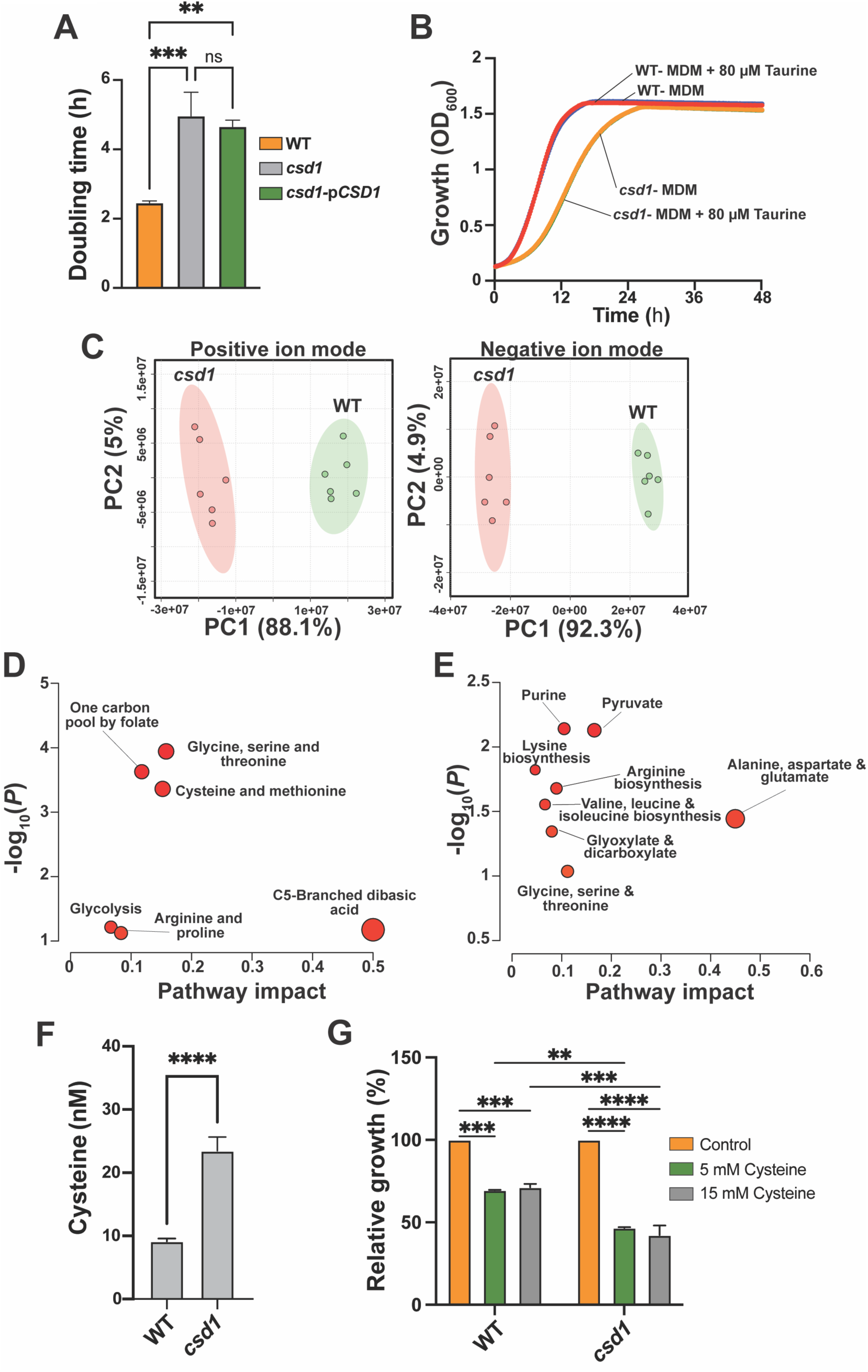
Genetic inactivation of *CSD1* disrupts sulfur amino acid homeostasis and impairs growth. (**A**) Doubling time of WT, *csd1* and *csd1*-p*CSD1* complemented cells grown in minimal dextrose medium (MDM). (**B**) Growth of WT and *csd1* cells in MDM in the presence or absence of 80 µM taurine. Exogenous taurine did not rescue the growth defect of *csd1*. Growth assays performed with additional taurine concentrations are shown in Figure S3. (**C**) PCA of untargeted metabolomic profiles from WT and *csd1* cells acquired in positive (left) and negative (right) ionization modes. Shaded ellipses indicate the clustering of biological replicates for each strain. (**D-E**) Pathway analysis of metabolites significantly enriched (**D**) or depleted (**E**) in *csd1* relative to WT using MetaboAnalyst 6.0. Pathways are plotted according to enrichment significance [-log_10_(P)] and pathway impact derived from pathway topology analysis. (**F**) Intracellular cysteine levels in WT and *csd1* cells. (**G**) Growth of WT and *csd1* cells in the absence or presence of 5 or 15 mM cysteine, expressed relative to growth under the corresponding untreated condition. Data in A and G are presented as means from, at least, three independent biological replicates. Statistical significance was determined using one-way ANOVA followed by Tukey’s multiple-comparisons test. NS, not significant; **P < 0.01; ***P < 0.001; ****P < 0.0001.

### *CSD1* promotes fungal virulence, adaptation to host-associated stresses and hyphal morphogenesis

The conservation of Csd1 in C. albicans and other pathogenic yeasts of the CTG clade, together with its absence in S. cerevisiae, prompted us to investigate whether Csd1 contributes to fungal fitness under host-associated conditions. To determine whether Csd1 contributes to fungal virulence *in vivo*, we used the *Galleria mellonella* model of systemic candidiasis. Infection with the WT strain resulted in marked larval mortality, with only 15% of larvae surviving 6 days post-infection (**Figure 3A**). In contrast, larvae injected with PBS or infected with the *csd1* mutant exhibited significantly higher survival rates, with 87% of larvae surviving at day 6. Infection with the *csd1*-p*CSD1* complemented strain led to an intermediate survival rate of 37%, partially restoring virulence toward WT levels. Collectively, these results support a specific contribution of Csd1 to *C. albicans* virulence in this model.

**Figure 3.**
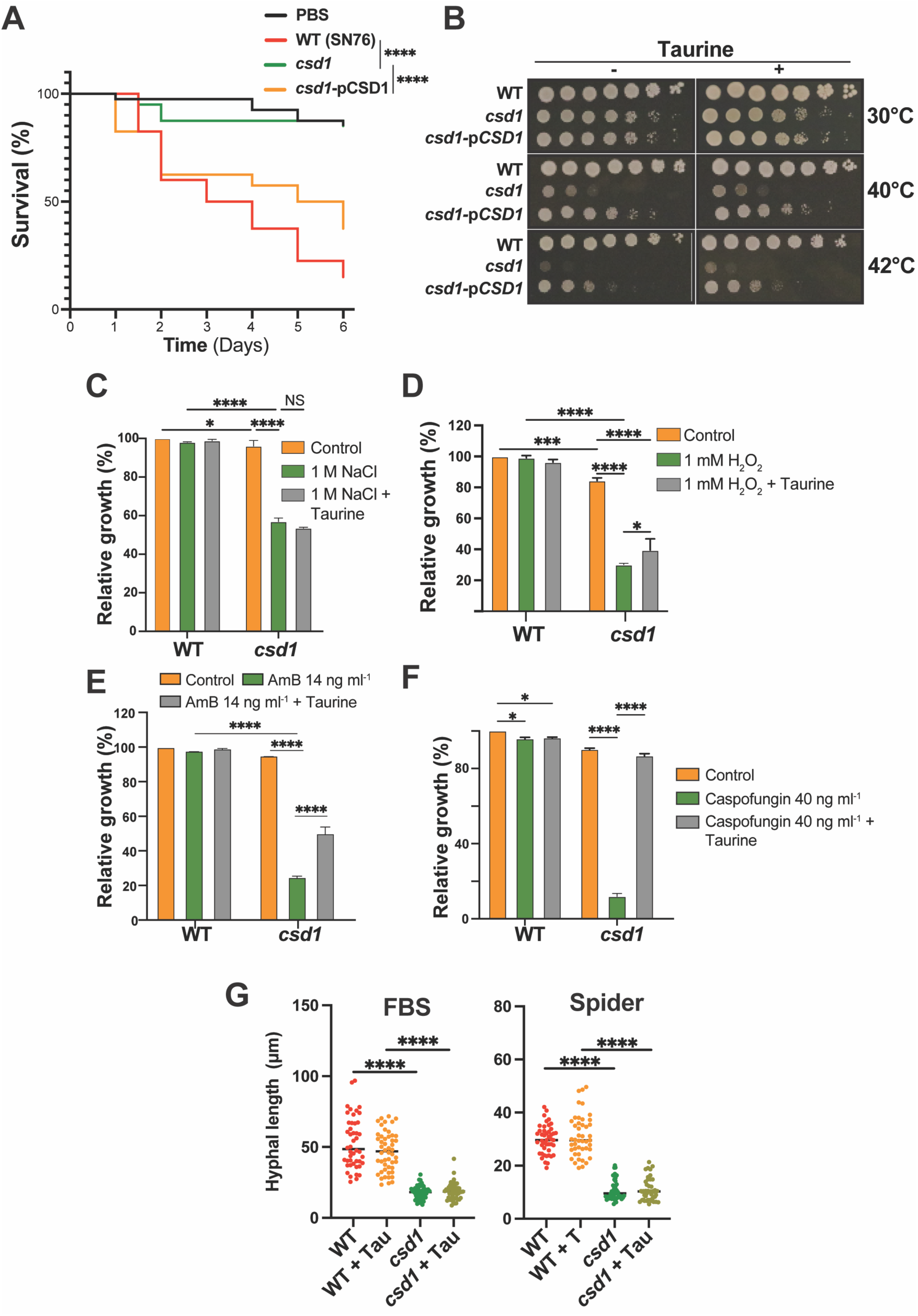
Loss of *CSD1* impairs virulence, stress tolerance, and hyphal development in *C. albicans*. (**A**) Kaplan-Meier survival analysis of *G. mellonella* larvae injected with PBS or infected with WT, *csd1*, or *csd1*-p*CSD1* complemented cells. (**B**) Serial-dilution growth assay of WT, *csd1*, and *csd1*-p*CSD1* cells at 30, 40, and 42°C in the presence or absence of 50 µM taurine. (**C-D**) Growth of WT and *csd1* cells under osmotic stress (**C**; 1 M NaCl) or oxidative stress (**D**; 1 mM H_2_O_2_), in the presence or absence of 50 µM taurine. Growth is expressed relative to the untreated WT condition. (**E-F**) Growth of WT and *csd1* cells in the presence of amphotericin B (**E**; AmB, 14 ng ml^−1^) or caspofungin (**F**; 40 ng ml^−1^), with or without 50 µM taurine. (**G**) Hyphal length of WT and *csd1* cells following induction with fetal bovine serum (FBS; left) or Spider medium (right), in the presence or absence of 50 µM taurine. Each point represents an individual hypha, and horizontal lines indicate the mean. Data in **C-F** are presented as means from at least three independent biological replicates. NS, not significant; *P < 0.05; ***P < 0.001; ****P < 0.0001. Survival curves in **A** were compared using the log-rank (Mantel-Cox) test.

Given the established roles of taurine in cellular stress responses across diverse organisms [11], we examined whether loss of *CSD1* altered the ability of *C. albicans* to withstand environmental and host-associated stresses summarized in **Table 1**. Phenotypic profiling revealed that the *csd1* mutant was particularly sensitive to thermal stress, displaying severe growth defects at both 40 and 42°C (**Figure 3B**). The mutant also exhibited pronounced hypersensitivity to osmotic stress imposed by 0.5-1 M NaCl (**Figure 3C**) and increased sensitivity to oxidative stress induced by H_2_O_2_ (**Figure 3D**). In addition, *csd1* cells showed increased sensitivity to sodium bisulfite, indicating an impaired ability to tolerate reactive sulfur species (**Table 1**). A mild growth defect was also observed under excess copper conditions (**Table 1**). Loss of *CSD1* also markedly increased susceptibility to antifungal drugs. The *csd1* mutant exhibited pronounced hypersensitivity to amphotericin B and caspofungin (**Figure 3E-F and Table 1**). Together, these phenotypes demonstrate that Csd1 contributes to the ability of *C. albicans* to withstand multiple physiologically relevant stresses, including thermal, oxidative, osmotic, reactive sulfur, and antifungal stresses.

**Table 1.**
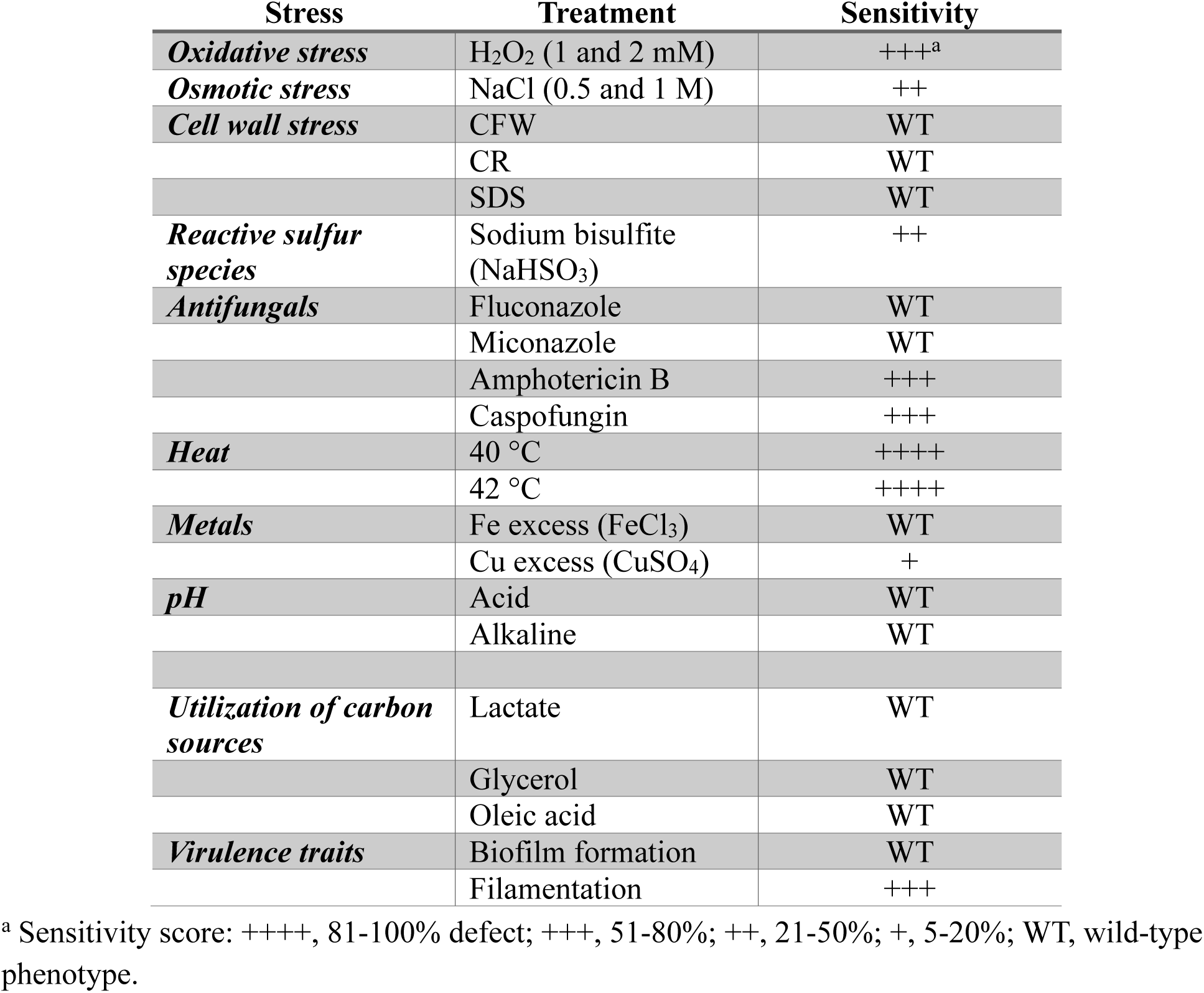
Phenotypic profile of the *C. albicans csd1* mutant under stress and host-relevant conditions.

We next asked whether exogenous taurine could alleviate the most pronounced stress phenotypes associated with *CSD1* loss. The extent of rescue was strongly condition dependent. Taurine fully restored growth in the presence of caspofungin (**Figure 3F**) and partially alleviated the defects associated with amphotericin B, elevated temperature, and oxidative stress (**Figure 3B and 3D-E**). In contrast, taurine supplementation failed to improve growth under osmotic stress, similar to its inability to rescue the intrinsic growth defect of the *csd1* mutant (**Figures 2B and 3C**). These differential responses suggest that some of the phenotypes associated with *CSD1* loss are linked to taurine deficiency, whereas others likely reflect broader metabolic perturbations resulting from disruption of taurine biosynthesis.

To determine whether Csd1 contributes to virulence-associated traits of *C. albicans*, we assessed its role in filamentation and biofilm formation. In response to fetal bovine serum (FBS), the *csd1* mutant formed significantly shorter hyphae (17.9 µm) than the WT (52.9 µm) (**Figure 3G**). A similar defect was observed in Spider medium, where *csd1* hyphae (10.7 µm) were markedly shorter than those of the WT (29.8 µm) (**Figure 3G**). Notably, exogenous taurine failed to restore hyphal elongation of the *csd1* mutant under either condition, indicating that the filamentation defect is not simply attributable to taurine deficiency. In contrast, loss of *CSD1* had no detectable effect on biofilm formation, with no significant difference between the *csd1* mutant and WT (**Table 1**). Together, these findings identify Csd1 as an important determinant of hyphal morphogenesis, while indicating that it is dispensable for biofilm formation under the conditions tested.

## Discussion

Our understanding of fungal sulfur metabolism has been shaped predominantly by studies of cysteine and methionine biosynthesis and uptake, together with the diverse metabolites derived from these sulfur-containing amino acids, such as the antioxidant glutathione and the principal cellular methyl donor SAM. Sulfur metabolism in human fungal pathogens also extends beyond these core pathways to specialized metabolites, including the mycotoxin gliotoxin and the antioxidant ergothioneine in *Aspergillus* species [33, 34]. Although taurine and hypotaurine have been detected in fungi such as *Y. lipolytica* and *G. candidum*, and components of the CSA pathway are conserved across several fungal species, the genetic basis and physiological significance of endogenous taurine production remain largely unresolved in fungi [25]. Here, we establish endogenous taurine production in *C. albicans*, identify Csd1 as its major genetic determinant, and show that Csd1-dependent metabolism is closely linked to sulfur amino acid homeostasis, stress adaptation, morphogenesis, and pathogenic fitness. Together, these findings reveal endogenous taurine production as a previously unrecognized facet of fungal sulfur metabolism.

Sequence conservation between Csd1 and characterized CSADs also points to an unexpected evolutionary relationship between fungal Csd1 and the enzymatic machinery underlying taurine biosynthesis in other organisms. Although CSAD-dependent taurine production is best characterized in mammals, *bona fide* CSAD activity has also been demonstrated in bacteria, including *Synechococcus*, indicating that PLP-dependent decarboxylation of cysteine sulfinic acid or cysteic acid is not restricted to metazoans [22]. Notably, Csd1 retains key features of the conserved CSAD catalytic machinery, including the PLP-binding lysine and conserved active-site residues, yet differs at residues implicated in substrate recognition. Studies of mammalian and bacterial CSADs have shown that relatively few residues within the substrate-binding pocket can profoundly influence discrimination among cysteine sulfinic acid, cysteic acid, and related acidic amino acids [22, 29, 30]. Thus, the divergent Y-X_19_-N-X-H signature of Csd1, compared with the canonical F/W-X_19_-S-X-Y motif of characterized CSADs, may reflect evolutionary remodeling of substrate recognition in the fungal enzyme. Whether this divergence alters substrate affinity or permits broader substrate promiscuity remains to be determined. Together, these findings suggest that Csd1 has retained key features of the conserved CSAD catalytic machinery while exhibiting a distinct predicted substrate-recognition architecture.

Our genetic analyses suggest that taurine biosynthesis in *C. albicans* differs from the canonical CSA pathway described in metazoans [11]. In this pathway, the cysteine dioxygenase Cdg1 catalyzes the conversion of cysteine to CSA, which is subsequently decarboxylated toward taurine. Unexpectedly, deletion of *CDG1*, encoding the predicted cysteine dioxygenase, did not reduce taurine levels but instead resulted in a modest increase, whereas loss of *GAD1* had no detectable effect. These findings indicate that neither Cdg1 nor Gad1 is individually required for taurine production under the conditions tested and suggest the existence of alternative or redundant routes feeding the pathway. In contrast, loss of *CSD1* reduced taurine to near the detection limit, establishing Csd1 as the major genetic determinant of taurine production, although residual synthesis through an alternative route cannot be excluded. Intriguingly, *csd1* cells retained the ability to utilize CSA as a sulfur source but failed to grow on cysteic acid, suggesting that CSA may represent a metabolic branch point that can enter a Csd1-independent route, whereas cysteic acid utilization requires Csd1. This raises the possibility that Csd1 preferentially participates in a cysteic acid-dependent route to taurine rather than functioning exclusively through the canonical CSA-to-hypotaurine route. Collectively, these observations support a branched and potentially redundant architecture for taurine biosynthesis in *C. albicans*, rather than simple conservation of the canonical metazoan pathway.

The pronounced growth defect of the *csd1* mutant provides an important clue to the physiological function of taurine biosynthesis. Loss of *CSD1* was accompanied by extensive remodeling of sulfur amino acid metabolism, including accumulation of cysteine and several cysteine- and methionine-associated metabolites, together with increased sensitivity to exogenous cysteine. Cysteine homeostasis is particularly important because excess cysteine can promote the production of sulfite, a reactive sulfur species that requires dedicated detoxification mechanisms in *C. albicans* [31, 35]. Consistent with a broader disruption of sulfur homeostasis, *csd1* cells were also hypersensitive to sulfite (**Figure S4**). Together, these phenotypes suggest that taurine biosynthesis may provide a metabolic sink for excess cysteine-derived sulfur, thereby limiting the accumulation or toxicity of reactive sulfur intermediates. This model is further supported by the high tolerance of *C. albicans* to exogenous taurine (**Figure S5**) and its capacity to mobilize taurine-derived sulfur through desulfonation [10, 36]. We therefore propose that taurine biosynthesis may function as a sulfur-buffering mechanism, channeling excess cysteine-derived sulfur into a well-tolerated metabolite that can subsequently be remobilized when sulfur becomes limiting. Such a mechanism could complement sulfite detoxification through Ssu1-mediated efflux [31], which protects against RSS toxicity but results in the loss of assimilated sulfur.

Beyond its potential role in sulfur homeostasis, our findings point to a broader contribution of taurine biosynthesis to cellular stress adaptation *in C. albicans*. Genetic inactivation of *CSD1* increased susceptibility to diverse environmental stresses, including oxidative and osmotic stress and elevated temperature, as well as to the antifungals caspofungin and amphotericin B. Importantly, exogenous taurine fully or partially restored several of these phenotypes, supporting a contribution of taurine deficiency to specific stress vulnerabilities, whereas the lack of rescue under other conditions suggests additional consequences of disrupting Csd1-dependent metabolism. These pleiotropic effects suggest that taurine may influence multiple aspects of fungal stress physiology, potentially involving redox homeostasis and membrane or cell-wall homeostasis. Interestingly, this phenotypic spectrum overlaps with that resulting from depletion of the thioredoxin reductase Trr1 [37], raising the possibility that perturbation of taurine and sulfur homeostasis intersects with cellular thiol-redox networks. Future studies will be required to determine whether these phenotypes reflect a direct role for taurine in cellular stress protection or secondary consequences of sulfur metabolic imbalance, and how these processes ultimately contribute to hyphal morphogenesis and virulence.

## Methods

### Strain construction and growth conditions

*C. albicans* strains and primers used for mutant construction are listed in **Table S2**. Unless otherwise indicated, strains were routinely grown and maintained at 30°C in yeast extract-peptone-dextrose (YPD) medium supplemented with uridine (2% Bacto peptone, 1% yeast extract, 2% [w/v] glucose, and 50 µg/ml uridine). Minimal dextrose medium (MDM; 1.7 g/l yeast nitrogen base without amino acids and ammonium sulfate, 5 g/l (NH_4_)_2_SO_4_, 2% [w/v] glucose, and 50 mg/l uridine) was used for phenotypic assays. Cysteic acid and cysteine sulfinic acid utilization assays were performed in sulfur-lacking dextrose (SLD) medium (1.2 g/l yeast nitrogen base without amino acids, ammonium sulfate, or magnesium, 4 g/l NH_4_Cl, 0.84 g/l MgCl_2_-6H_2_O, and 20 g/l glucose), supplemented with cysteic acid or cysteine sulfinic acid as the sole sulfur source.

The *csd1* deletion mutant was generated in the SN148 background by replacing the entire *CSD1* open reading frame (ORF) with PCR-amplified disruption cassettes derived from pFA plasmids, as previously described [38]. For complementation, the *CSD1* ORF was amplified and cloned downstream of the *ACT1* promoter in the CIp-ACT integrating vector [39]. The resulting construct was linearized with *StuI* and integrated at the *RPS1* locus of the *csd1* mutant using lithium acetate transformation [40]. Transformants were selected on synthetic complete medium (SC) lacking uridine (0.67% yeast nitrogen base without amino acids, 2% glucose, and 0.08% dropout mix supplemented with the required amino acids), and correct integration was verified by PCR.

### Phenotypic assays

For liquid growth assays, overnight cultures of the different *C. albicans* strains grown in SC medium were diluted to an OD_600_ of 0.1 and dispensed into flat-bottom 96-well plates at a final volume of 200 µl per well in the presence of the indicated antifungals or stressors. Each plate included a drug- or stressor-free growth control and a cell-free negative control. Growth was monitored over 48h at 30°C using a Sunrise™ microplate reader (Tecan) under continuous agitation. Antifungal agents were dissolved in dimethyl sulfoxide (DMSO) and included fluconazole (128 mg/ml; Cayman Chemical, catalog no. 11594-5), miconazole (8 mg/ml; Sigma, catalog no. M3512), caspofungin (8 mg/ml; Merck, Cancidas; catalog no. 02244266), and amphotericin B (1 mg/ml; BioBasic, catalog no. AD0030P). Sulfite tolerance assays were performed using overnight cultures grown in SC medium adjusted to pH 3.5. Cells were exposed to the indicated concentrations of freshly prepared sodium bisulfite (NaHSO_3_), and growth was monitored at 30 °C using a Cytation 5 plate reader under continuous agitation. OD_600_ measurements were recorded every 10 min for 48 h.

Heat-stress assays were performed on solid medium. Overnight cultures grown in SC medium were adjusted to an OD_600_ of 4 and subjected to serial 1:50 dilutions. Two microliters of each dilution were spotted onto SC agar plates and incubated for 48 h at 30, 40, or 42°C. Images were captured using the SP-imager system (S&P Robotics, Toronto, ON, Canada).

Hyphal induction was assessed in liquid Spider medium and YPD supplemented with 10% FBS. Overnight cultures grown in YPD at 30°C were washed with PBS, diluted to an OD_600_ of 0.05 in pre-warmed medium, and incubated at 37°C for 3 h to induce filamentation. Cells were imaged by bright-field microscopy (Nikon Instruments Inc., Melville, NY, USA) using a 40× objective. Hyphal length was measured for at least 45 cells per strain and condition using NIS-Elements imaging software. Measurements were obtained from three independent biological replicates. Biofilm formation was performed as described previously [41].

### Taurine quantification

C. albicans strains were inoculated into 50 ml of MDM medium at an initial OD_600_ of 0.2 and grown at 30°C to logarithmic phase (OD_600_ = 0.8). Cells were harvested by centrifugation at 13,000 × g for 5 min, washed with cold PBS, and transferred to pre-chilled 1.5-ml tubes. Cell pellets were resuspended in 700 µl of ice-cold extraction buffer consisting of acetonitrile, formic acid, and deionized water (10:0.1:90, v/v/v), and 100 µl of 0.1-mm zirconia beads (BioSpec) were added.

Cells were disrupted using a Mini-Beadbeater-24 (BioSpec) at 6,800 rpm for eight 30s cycles, with 45s intervals at 4°C between cycles. Lysates were centrifuged at 10,000 rpm for 10 min at 4°C, and the resulting supernatants were collected and stored at −80°C until analysis. A total of five independent biological replicates were analyzed.

Chromatographic separation was performed at the Centre for Structural and Functional Genomics (Concordia University, Montreal, QC) using an Agilent 1290 Infinity II ultrahigh-performance liquid chromatography system (Agilent Technologies, Santa Clara, CA, USA). Fifteen microliters of each sample were injected onto a Luna NH_2_ column (2.1 × 250 mm, 5 µm; Phenomenex). The mobile phases consisted of 0.1% formic acid in water (solvent A) and 0.1% formic acid in acetonitrile (solvent B). The gradient was increased from 3% to 20% B over 3 min, followed by an increase from 20% to 85% B over 1 min and a return to 3% B over 0.1 min. The column was then re-equilibrated at 3% B for 4 min. The flow rate was 350 µl/min. The column was connected in-line to an Agilent 6560 ion mobility quadrupole time-of-flight (QTOF) mass spectrometer equipped with a dual Jet Stream electrospray ionization source. Data were acquired in negative-ion mode over an m/z range of 50-1100 at an acquisition rate of 2 Hz. Taurine was quantified using MassHunter Quantitative Analysis software (Agilent Technologies), with a 5-ppm extraction window centered on the taurine target ion (m/z 124.0074) for peak-area determination.

### Untargeted metabolomics

#### Metabolite extraction and derivatization

Approximately 200 mg of pre-washed yeast cell pellets were transferred to 2 ml tubes containing approximately 200 mg of 0.5 mm glass beads. Metabolites were extracted using a cold MeOH:ACN:H_2_O mixture (2:2:1, v/v/v) at a ratio of 6 µl of extraction solvent per mg of yeast. Samples were homogenized by bead beating (BeadRuptor, Omni International, Kennesaw, GA, USA) and centrifuged at 14,000 rpm for 8 min at 4°C. The resulting supernatants were collected for metabolomic analysis, with 600 µl used for underivatized analysis and 200 µl for phenylisothiocyanate (PITC) derivatization. Aliquots for underivatized analysis were dried and reconstituted in 300 µl of 25% MeOH before LC-MS/MS analysis. For PITC derivatization, dried extracts were resuspended in 200 µl of EtOH:H₂O:pyridine (1:1:1, v/v/v) containing 0.4 M PITC and incubated for 1 h at room temperature with gentle mixing. Samples were subsequently dried and reconstituted in 100 µl of 25% MeOH before LC-MS/MS analysis.

#### LC-MS/MS analysis

Untargeted metabolomic profiling was performed using a Shimadzu Nexera UHPLC system coupled to a SCIEX X500B quadrupole time-of-flight (QqTOF) mass spectrometer (SCIEX, Concord, ON, Canada). Chromatographic separation was achieved using water containing 0.1% formic acid as mobile phase A and acetonitrile containing 0.1% formic acid as mobile phase B, under gradient-elution conditions. Samples were analyzed independently in positive- and negative-ion electrospray ionization modes (Scherzo^+^ and Scherzo^−^, respectively) to maximize metabolite coverage. Data were acquired using high-resolution time-of-flight mass spectrometry using data-dependent acquisition. Full-scan TOF-MS spectra were acquired over m/z 100-1,000 with a 250 ms accumulation time, and MS/MS spectra were acquired over m/z 40-900 with a 150 ms accumulation time. Collision energy was set to 30 ± 10 V, and MS/MS spectra were acquired for up to the eight most intense precursor ions per cycle.

#### Data processing and metabolite identification

LC-MS features were generated on the basis of accurate mass-to-charge ratio (m/z) and retention time. Positive- and negative-ion datasets were processed independently. Metabolites were identified by comparison of experimental MS/MS spectra with a reference spectral library, yielding 90 identified metabolites in negative-ion mode and 112 in positive-ion mode. After accounting for metabolites detected in both ionization modes, 163 unique metabolites were identified. Metabolite abundances in *csd1* and WT samples were compared using integrated chromatographic peak areas from the identified metabolites. Positive- and negative-ion datasets were initially evaluated separately, and PCA was used to assess global differences in metabolic profiles between WT and *csd1* samples. Differential metabolites were identified based on statistical significance and fold change [1.5 fold-change and FDR 5%]. Metabolites detected in both ionization modes were consolidated to generate a non-redundant set for downstream interpretation (**Table S1**). Pathway analysis was performed using MetaboAnalyst 6.0 [42]. Differentially abundant metabolites were mapped to metabolic pathways, and pathway significance was evaluated using enrichment analysis together with pathway-topology analysis.

### Cysteine quantification

Intracellular cysteine was quantified using the Cysteine Colorimetric Assay Kit (Thermo Fisher Scientific, Cat. No. EEA045) according to the manufacturer’s instructions. Fungal cells were grown to exponential phase in 50 ml of MDM medium at 30°C with shaking at 200 rpm. Cells were washed twice with PBS and resuspended in 200 µl of the kit-supplied lysis buffer with 100 µl of 0.1-mm glass beads. Cells were lysed using a BioSpec Mini-Beadbeater-24 at 6,800 rpm, and lysates were clarified by centrifugation at 10,000 x g for 10 min at 4°C. The resulting supernatants were collected and used for cysteine quantification. Cysteine levels were normalized to cell number and expressed as nmol per 10^6^ cells. Measurements were performed using four independent biological replicates.

### Galleria mellonella infection assay

Virulence assays were performed using *G. mellonella* larvae obtained from Elevages Lisard (Saint-Bruno-de-Montarville, QC, Canada), as previously described [43]. Larvae were randomly assigned to groups of 20, and individuals displaying signs of melanization before infection were excluded. Overnight cultures of *C. albicans* strains grown in YPD medium were washed twice and resuspended in PBS to a final inoculum of 5 × 10^5^ cells in 20 μl. Larvae were injected into the last left proleg, while control larvae were injected with an equivalent volume of sterile PBS. Larvae were incubated at 37 °C, and mortality was recorded daily for 6 days. Larvae were considered dead when they failed to respond to touch and were unable to right themselves. Two independent experiments, each comprising 20 larvae per condition, were performed. Kaplan–Meier survival curves were generated and compared using the log-rank test (GraphPad Prism 10).

### Statistical analyses

Statistical analyses were performed using GraphPad Prism 10. Unless otherwise indicated, data were obtained from at least three independent biological replicates and are presented as means ± standard deviation (SD). Statistical significance among multiple groups was assessed using one-way analysis of variance (ANOVA) followed by Tukey’s multiple-comparisons test. Survival curves were analyzed using the Kaplan–Meier method and compared using the log-rank test. Statistical significance was defined as follows: *P < 0.05, **P < 0.01, ***P < 0.001, and ****P < 0.0001.

## Acknowledgments

We thank all members of Dr. Sellam’s lab for their valuable comments and discussions. We also thank Dr Marcos Di Falco and Mindy Melgar (Concordia University, Montreal, QC) for their assistance in optimizing the LC-MS/MS protocol for taurine quantification. This work was supported by a Canadian Institutes for Health Research project grant (grant PJT-180256), the Canada Foundation for Innovation and the Montreal Heart Institute foundation to Adnane Sellam. Anagha C.T. Menon was supported by a CREATE doctoral scholarship from the Natural Sciences and Engineering Research Council of Canada (NSERC) through the EvoFunPath program. Adnane Sellam is supported by a Fonds de Recherche du Québec-Santé Senior Salary award.

## Supplementary Figures

**Figure S1.**
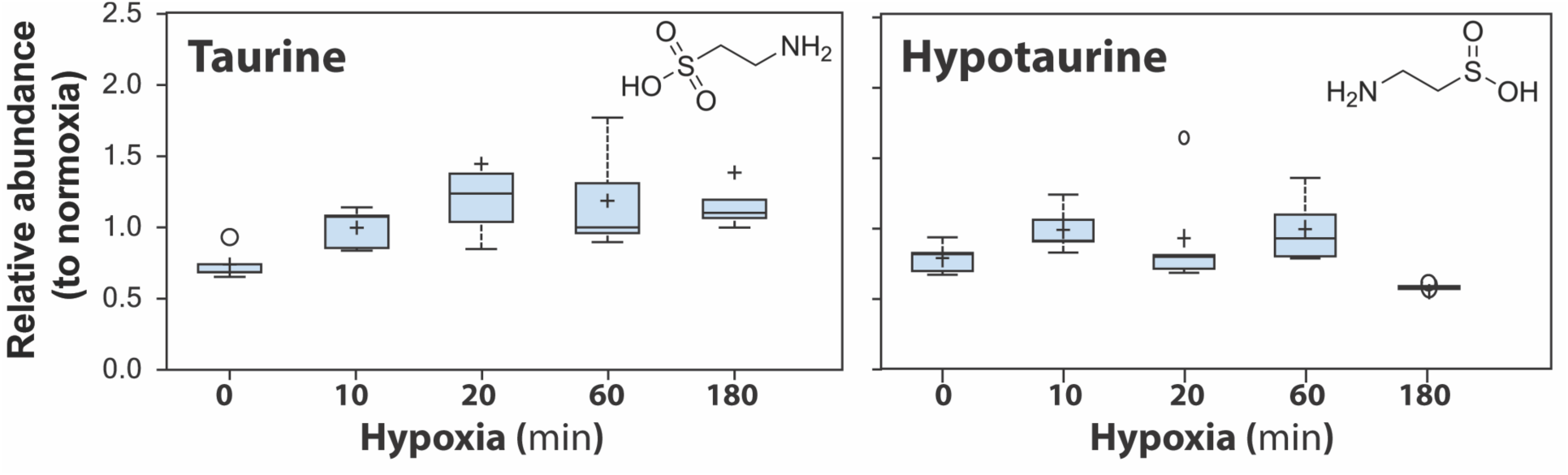
Relative abundance of taurine and hypotaurine in C. *albicans* cells exposed to hypoxia. Fungal cells were grown in YPD medium and exposed to hypoxia (5% O_2_) for 10, 20, 60 and 180 minutes. Taurine was detected by ultra high-performance liquid chromatography-tandem mass spectrometry (Metabolon, Durham, NC, USA). Results represent the mean of taurine levels of five biological replicates under hypoxia relative to normoxia.

**Figure S2.**
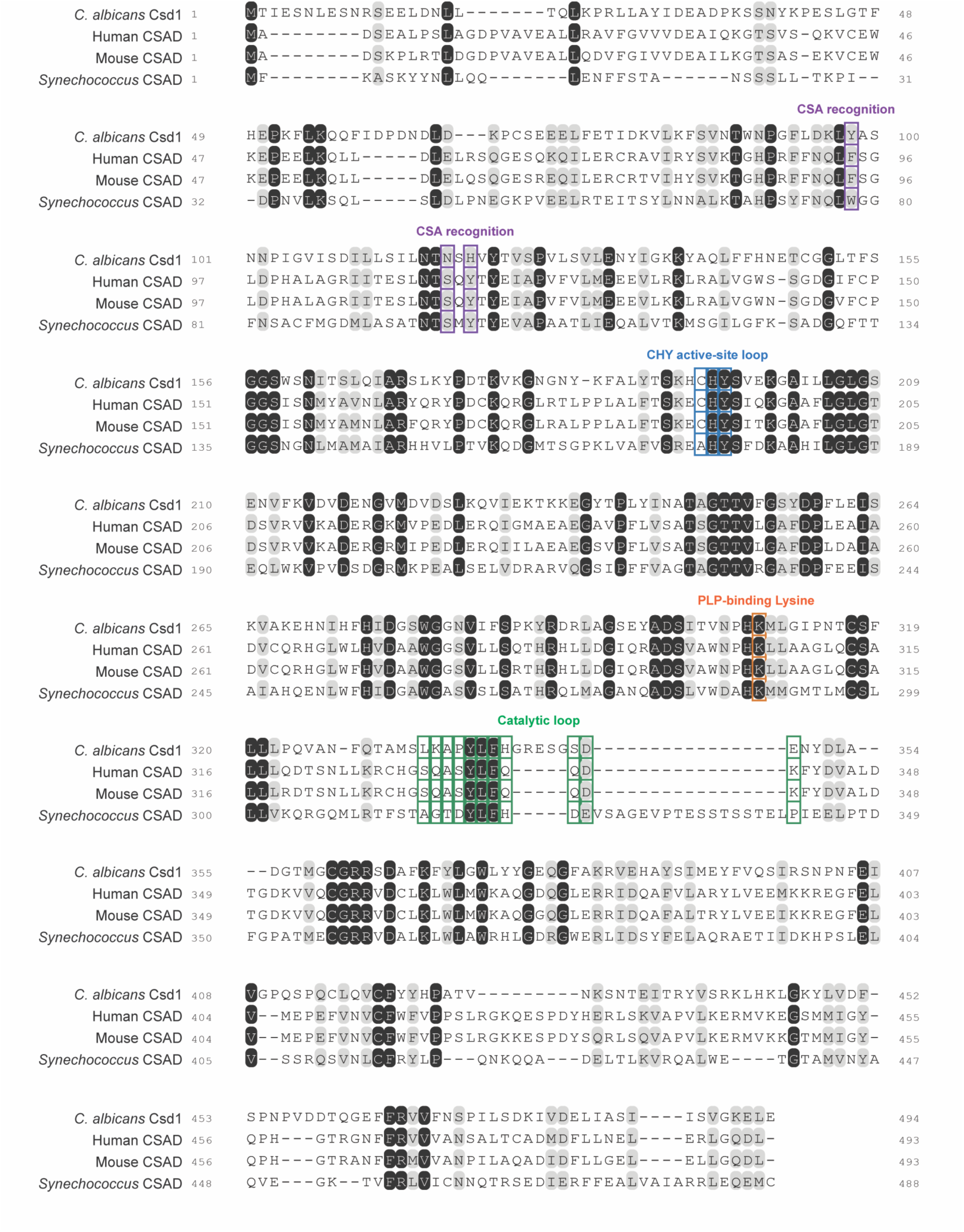
Sequence alignment of fungal, mammalian and bacterial cysteine sulfinate decarboxylases. Multiple sequence alignment of C. *albicans* Csdl, human CSAD, mouse CSAD, and the experimentally characterized CSAD from *Synechococcus sp.* PCC 7335. Dark shading indicates amino acid residues identical across all four proteins, whereas light shading indicates conservative substitutions. Key functional features of CSAD are annotated, including the cysteine sulfinic acid and the cysteic acid recognition residues (F94, S114, and Y116), the active-site loop (C190-H191-Y192), the PLP-binding lysine (K305), and the putative catalytic loop (S331-K341). Functional annotations are based on human CSAD numbering, with the corresponding aligned residues shown for the other proteins.

**Figure S3.**
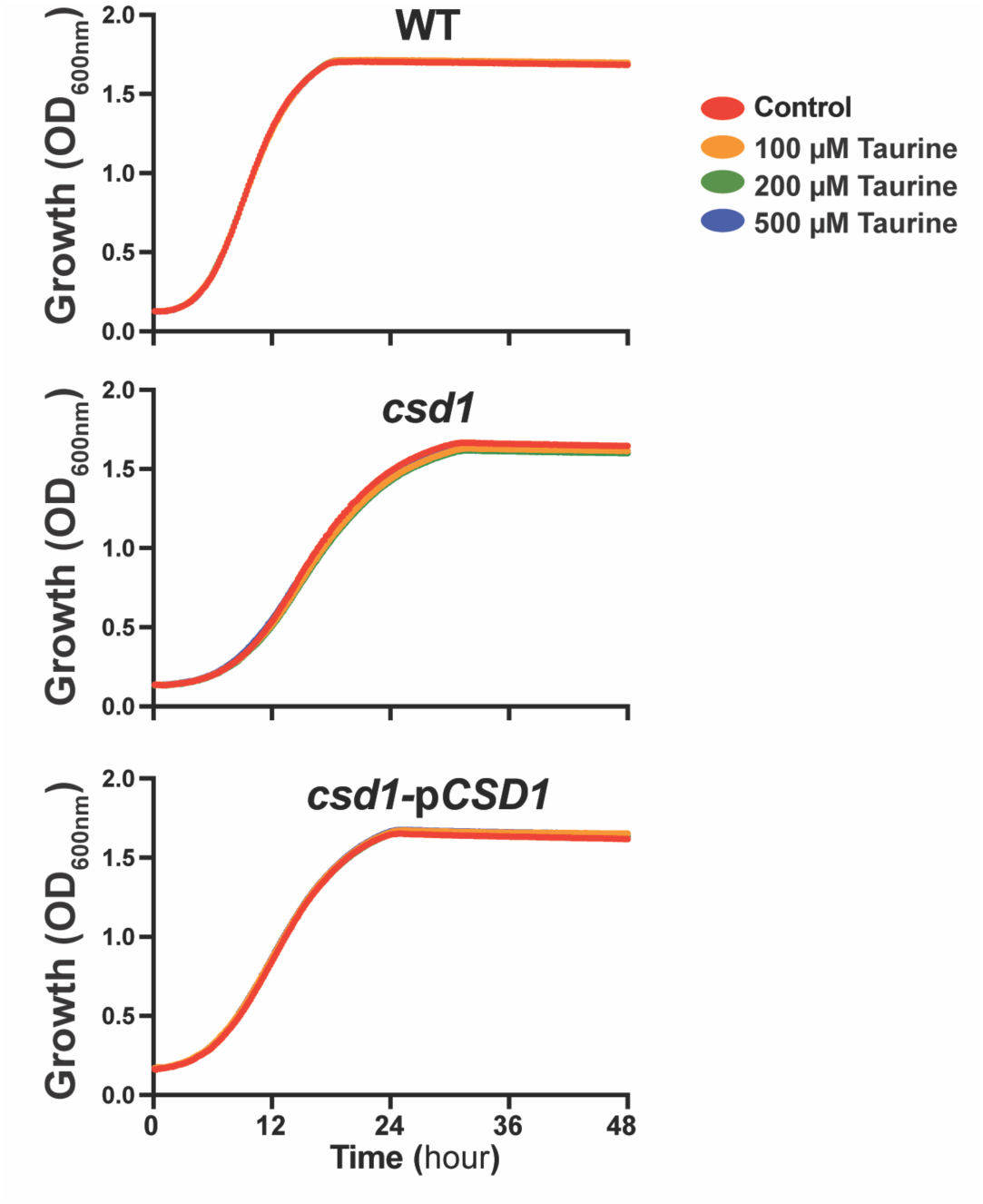
Effect of exogenous taurine on growth of the *csd1* mutant. Growth curves of WT, *csd1,* and csc/7-pCSD1 complemented strains in minimal dextrose medium (MDM) supplemented with the indicated concentrations of taurine. Exogenous taurine did not rescue the intrinsic growth defect of the *csd1* mutant, consistent with the analysis shown in **Figure 2B.**

**Figure S4.**
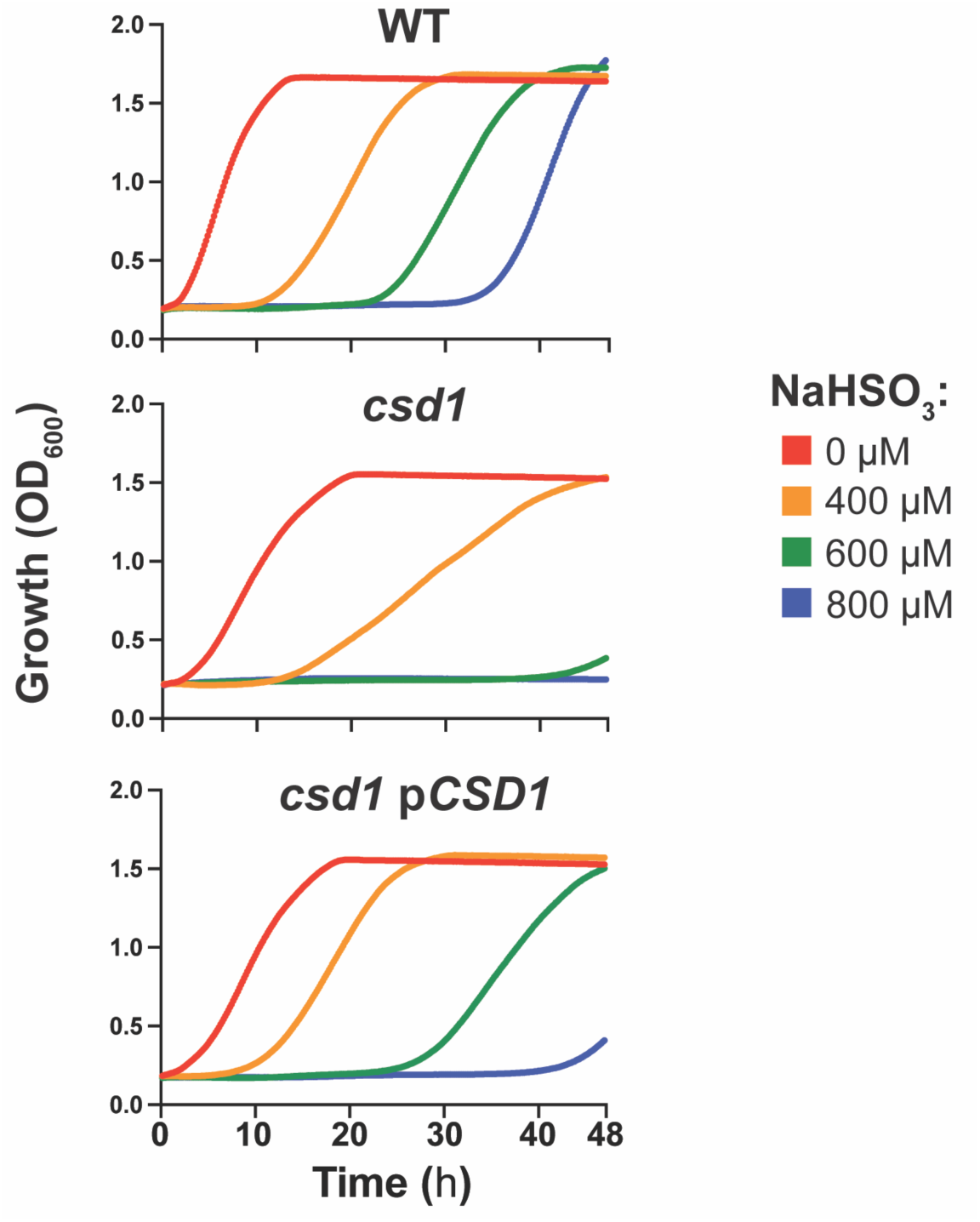
Loss of *CSD1* increases sensitivity to sulfite. Growth curves of WT, *csd1,* and *csd1-pCSD1* complemented strains in minimal dextrose medium (MDM) supplemented with the indicated concentrations of sodium bisulfite (NaHSO_3_). Growth was monitored by measuring OD_600_ over 48 h.

**Figure S5.**
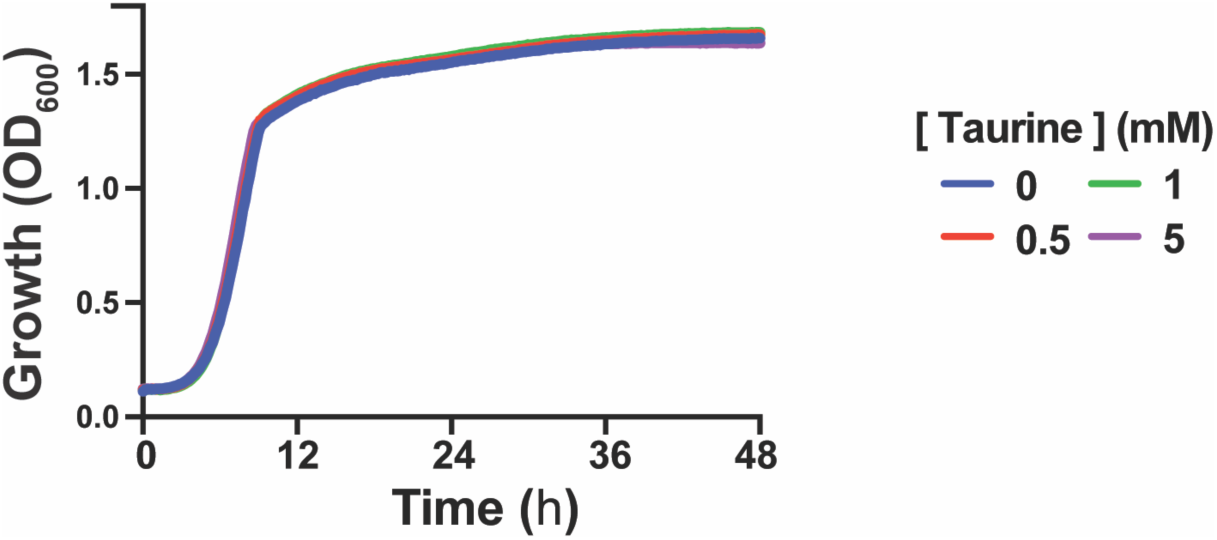
Effect of exogenous taurine on growth of C. *albicans.* Growth curves of the WT strain (SN250) in minimal dextrose medium (MDM) supplemented with the indicated concentrations of taurine. High concentrations of exogenous taurine did not impair C. *albicans* growth.

**Table S1.** Differential metabolomic profiling of WT and *csd1* mutant cells.

**Table S2.** Primers and strains used in this study.

## References

1. Brown GD, Denning DW, Gow NA, Levitz SM, Netea MG, White TC. Hidden killers: human fungal infections. Sci Transl Med. 2012;4(165):165rv13. doi: 10.1126/scitranslmed.3004404. PubMed PMID: 23253612.

2. Denning DW. Global incidence and mortality of severe fungal disease. Lancet Infect Dis. 2024;24(7):e428–e38. Epub 20240112. doi: 10.1016/S1473-3099(23)00692-8. PubMed PMID: 38224705.

3. Pappas PG, Kauffman CA, Andes D, Benjamin DK, Jr., Calandra TF, Edwards JE, Jr., et al. Clinical practice guidelines for the management of candidiasis: 2009 update by the Infectious Diseases Society of America. Clin Infect Dis. 2009;48(5):503–35. doi: 10.1086/596757. PubMed PMID: 19191635; PubMed Central PMCID: PMCPMC7294538.

4. Pfaller MA, Diekema DJ. Epidemiology of invasive candidiasis: a persistent public health problem. Clin Microbiol Rev. 2007;20(1):133–63. doi: 10.1128/CMR.00029-06. PubMed PMID: 17223626; PubMed Central PMCID: PMCPMC1797637.

5. (WHO) WHO. WHO fungal priority pathogens list to guide research, development and public health action. World Health Organization (WHO), 2022.

6. Oliva A, De Rosa FG, Mikulska M, Pea F, Sanguinetti M, Tascini C, et al. Invasive Candida infection: epidemiology, clinical and therapeutic aspects of an evolving disease and the role of rezafungin. Expert Rev Anti Infect Ther. 2023;21(9):957–75. Epub 20230817. doi: 10.1080/14787210.2023.2240956. PubMed PMID: 37494128.

7. WHO TEAM ARA, Health Systems, Access and Data (HSD). Blueprint for strengthening responses to fungal disease and antifungal resistance. 2026.

8. Puumala E, Fallah S, Robbins N, Cowen LE. Advancements and challenges in antifungal therapeutic development. Clin Microbiol Rev. 2024;37(1):e0014223. Epub 20240131. doi: 10.1128/cmr.00142-23. PubMed PMID: 38294218; PubMed Central PMCID: PMCPMC10938895.

9. Lensmire JM, Hammer ND. Nutrient sulfur acquisition strategies employed by bacterial pathogens. Curr Opin Microbiol. 2019;47:52–8. Epub 20181207. doi: 10.1016/j.mib.2018.11.002. PubMed PMID: 30530037.

10. Menon ACT, Tebbji F, Mecteau M, Alikashani A, Vincent AT, Rheaume E, et al. Met32 governs transcriptional control of sulfur metabolic flexibility and resistance to reactive sulfur species in the human fungal pathogen Candida albicans. mBio. 2026;17(5):e0047226. Epub 20260420. doi: 10.1128/mbio.00472-26. PubMed PMID: 42003610; PubMed Central PMCID: PMCPMC13170222.

11. Huxtable RJ. Physiological actions of taurine. Physiol Rev. 1992;72(1):101–63. doi: 10.1152/physrev.1992.72.1.101. PubMed PMID: 1731369.

12. Wang L, Xie Z, Wu M, Chen Y, Wang X, Li X, et al. The role of taurine through endoplasmic reticulum in physiology and pathology. Biochem Pharmacol. 2024;226:116386. Epub 20240622. doi: 10.1016/j.bcp.2024.116386. PubMed PMID: 38909788.

13. Miyazaki T. Identification of a novel enzyme and the regulation of key enzymes in mammalian taurine synthesis. J Pharmacol Sci. 2024;154(1):9–17. Epub 20231121. doi: 10.1016/j.jphs.2023.11.003. PubMed PMID: 38081683.

14. Wishart DS, Guo A, Oler E, Wang F, Anjum A, Peters H, et al. HMDB 5.0: the Human Metabolome Database for 2022. Nucleic Acids Res. 2022;50(D1):D622–D31. doi: 10.1093/nar/gkab1062. PubMed PMID: 34986597; PubMed Central PMCID: PMCPMC8728138.

15. Linder T. Assimilation of alternative sulfur sources in fungi. World J Microbiol Biotechnol. 2018;34(4):51. Epub 20180317. doi: 10.1007/s11274-018-2435-6. PubMed PMID: 29550883; PubMed Central PMCID: PMCPMC5857272.

16. Stipanuk MH. Sulfur amino acid metabolism: pathways for production and removal of homocysteine and cysteine. Annu Rev Nutr. 2004;24:539–77. doi: 10.1146/annurev.nutr.24.012003.132418. PubMed PMID: 15189131.

17. Tevatia R, Allen J, Rudrappa D, White D, Clemente TE, Cerutti H, et al. The taurine biosynthetic pathway of microalgae. Algal Research. 2015;9:21–6. doi: 10.1016/j.algal.2015.02.012.

18. Joseph CA, Maroney MJ. Cysteine dioxygenase: structure and mechanism. Chem Commun (Camb). 2007;(32):3338–49. doi: 10.1039/b702158e. PubMed PMID: 18019494.

19. Liu P, Ge X, Ding H, Jiang H, Christensen BM, Li J. Role of glutamate decarboxylase-like protein 1 (GADL1) in taurine biosynthesis. J Biol Chem. 2012;287(49):40898–906. Epub 20121004. doi: 10.1074/jbc.M112.393728. PubMed PMID: 23038267; PubMed Central PMCID: PMCPMC3510794.

20. Nishikawa M. Identification of the Yarrowia lipolytica cysteine sulfinic acid decarboxylase gene using a newly developed method with optimized Escherichia coli combinations of mutant alleles. Microbiology (Reading). 2025;171(11). doi: 10.1099/mic.0.001620. PubMed PMID: 41187071; PubMed Central PMCID: PMCPMC12585060.

21. Edgar SE, Hickman MA, Marsden MM, Morris JG, Rogers QR. Dietary cysteic acid serves as a precursor of taurine for cats. J Nutr. 1994;124(1):103–9. doi: 10.1093/jn/124.1.103. PubMed PMID: 8283286.

22. Agnello G, Chang LL, Lamb CM, Georgiou G, Stone EM. Discovery of a substrate selectivity motif in amino acid decarboxylases unveils a taurine biosynthesis pathway in prokaryotes. ACS Chem Biol. 2013;8(10):2264–71. Epub 20130823. doi: 10.1021/cb400335k. PubMed PMID: 23972067; PubMed Central PMCID: PMCPMC3815685.

23. North JA, Shafaat HS. Function, Structure, and Regulation of Nitrogen Fixation-like Metalloproteins for Nitrogen, Energy, Carbon, and Sulfur Metabolism. Chem Rev. 2026;126(15):8234–83. doi: 10.1021/acs.chemrev.5c00922. PubMed PMID: 42503660; PubMed Central PMCID: PMCPMC13474568.

24. Pracharova P, Lieben P, Pollet B, Beckerich JM, Bonnarme P, Landaud S, et al. Geotrichum candidum gene expression and metabolite accumulation inside the cells reflect the strain oxidative stress sensitivity and ability to produce flavour compounds. FEMS Yeast Res. 2019;19(1). doi: 10.1093/femsyr/foy111. PubMed PMID: 30295727; PubMed Central PMCID: PMCPMC6211236.

25. Hebert A, Forquin-Gomez MP, Roux A, Aubert J, Junot C, Heilier JF, et al. New insights into sulfur metabolism in yeasts as revealed by studies of Yarrowia lipolytica. Appl Environ Microbiol. 2013;79(4):1200–11. Epub 20121207. doi: 10.1128/AEM.03259-12. PubMed PMID: 23220962; PubMed Central PMCID: PMCPMC3568587.

26. Burgain A, Tebbji F, Khemiri I, Sellam A. Metabolic Reprogramming in the Opportunistic Yeast Candida albicans in Response to Hypoxia. mSphere. 2020;5(1). Epub 20200226. doi: 10.1128/mSphere.00913-19. PubMed PMID: 32102943; PubMed Central PMCID: PMCPMC7045390.

27. Paietta JV. 12 Regulation of Sulfur Metabolism in Filamentous Fungi. In: Hoffmeister D, editor. Biochemistry and Molecular Biology. Cham: Springer International Publishing; 2016. p. 305–19.

28. Cook AM, Laue H, Junker F. Microbial desulfonation. FEMS Microbiol Rev. 1998;22(5):399–419. doi: 10.1111/j.1574-6976.1998.tb00378.x. PubMed PMID: 9990724.

29. Tramonti A, Contestabile R, Florio R, Nardella C, Barile A, Di Salvo ML. A Novel, Easy Assay Method for Human Cysteine Sulfinic Acid Decarboxylase. Life (Basel). 2021;11(5). Epub 20210514. doi: 10.3390/life11050438. PubMed PMID: 34068845; PubMed Central PMCID: PMCPMC8153620.

30. Mahootchi E, Raasakka A, Luan W, Muruganandam G, Loris R, Haavik J, et al. Structure and substrate specificity determinants of the taurine biosynthetic enzyme cysteine sulphinic acid decarboxylase. J Struct Biol. 2021;213(1):107674. Epub 20201127. doi: 10.1016/j.jsb.2020.107674. PubMed PMID: 33253877.

31. Hennicke F, Grumbt M, Lermann U, Ueberschaar N, Palige K, Bottcher B, et al. Factors supporting cysteine tolerance and sulfite production in Candida albicans. Eukaryot Cell. 2013;12(4):604–13. Epub 20130215. doi: 10.1128/EC.00336-12. PubMed PMID: 23417561; PubMed Central PMCID: PMCPMC3623443.

32. Wain WH, Price MF, Cawson RA. A re-evaluation of the effect of cysteine or Candida albicans. Sabouraudia. 1975;13 Pt 1:74–82. PubMed PMID: 1092000.

33. Traynor AM, Sheridan KJ, Jones GW, Calera JA, Doyle S. Involvement of Sulfur in the Biosynthesis of Essential Metabolites in Pathogenic Fungi of Animals, Particularly Aspergillus spp.: Molecular and Therapeutic Implications. Front Microbiol. 2019;10:2859. Epub 20191213. doi: 10.3389/fmicb.2019.02859. PubMed PMID: 31921039; PubMed Central PMCID: PMCPMC6923255.

34. Amich J. The many roles of sulfur in the fungal-host interaction. Curr Opin Microbiol. 2024;79:102489. Epub 20240515. doi: 10.1016/j.mib.2024.102489. PubMed PMID: 38754292.

35. Chebaro Y, Lorenz M, Fa A, Zheng R, Gustin M. Adaptation of Candida albicans to Reactive Sulfur Species. Genetics. 2017;206(1):151–62. Epub 20170224. doi: 10.1534/genetics.116.199679. PubMed PMID: 28235888; PubMed Central PMCID: PMCPMC5419466.

36. Linder T. Genomics of alternative sulfur utilization in ascomycetous yeasts. Microbiology (Reading). 2012;158(Pt 10):2585–97. Epub 20120712. doi: 10.1099/mic.0.060285-0. PubMed PMID: 22790398.

37. Qi W, Roy U, Cai C, Acosta-Zaldivar M, Mascio J, Asara JM, et al. Cascading damage to Candida albicans cells through thioredoxin reductase loss. Proc Natl Acad Sci U S A. 2026;123(33):e2615451123. Epub 20260810. doi: 10.1073/pnas.2615451123. PubMed PMID: 42574622.

38. Gola S, Martin R, Walther A, Dunkler A, Wendland J. New modules for PCR-based gene targeting in Candida albicans: rapid and efficient gene targeting using 100 bp of flanking homology region. Yeast. 2003;20(16):1339–47. doi: 10.1002/yea.1044. PubMed PMID: 14663826.

39. Blackwell C, Russell CL, Argimon S, Brown AJ, Brown JD. Protein A-tagging for purification of native macromolecular complexes from Candida albicans. Yeast. 2003;20(15):1235–41. doi: 10.1002/yea.1036. PubMed PMID: 14618561.

40. Wilson RB, Davis D, Enloe BM, Mitchell AP. A recyclable Candida albicans URA3 cassette for PCR product-directed gene disruptions. Yeast. 2000;16(1):65–70. doi: 10.1002/(SICI)1097-0061(20000115)16:1<65::AID-YEA508>3.0.CO;2-M. PubMed PMID: 10620776.

41. Garcia C, Burgain A, Chaillot J, Pic E, Khemiri I, Sellam A. A phenotypic small-molecule screen identifies halogenated salicylanilides as inhibitors of fungal morphogenesis, biofilm formation and host cell invasion. Sci Rep. 2018;8(1):11559. Epub 20180801. doi: 10.1038/s41598-018-29973-8. PubMed PMID: 30068935; PubMed Central PMCID: PMCPMC6070544.

42. Pang Z, Lu Y, Zhou G, Hui F, Xu L, Viau C, et al. MetaboAnalyst 6.0: towards a unified platform for metabolomics data processing, analysis and interpretation. Nucleic Acids Res. 2024;52(W1):W398–W406. doi: 10.1093/nar/gkae253. PubMed PMID: 38587201; PubMed Central PMCID: PMCPMC11223798.

43. Burgain A, Pic E, Markey L, Tebbji F, Kumamoto CA, Sellam A. A novel genetic circuitry governing hypoxic metabolic flexibility, commensalism and virulence in the fungal pathogen Candida albicans. PLoS Pathog. 2019;15(12):e1007823. Epub 20191206. doi: 10.1371/journal.ppat.1007823. PubMed PMID: 31809527; PubMed Central PMCID: PMCPMC6919631.

